## supplementary figures for "The gut microbiome-linked long chain fatty acid stearate suppresses colorectal cancer"

### Extended data figure 1

**a**

| Donor number | 1 | 2 | 3 | 4 | 5 |
|---|---|---|---|---|---|
| Gender | F | F | M | F | M |
| Age (years) | 60 | 62 | 43 | 58 | 57 |

**b**

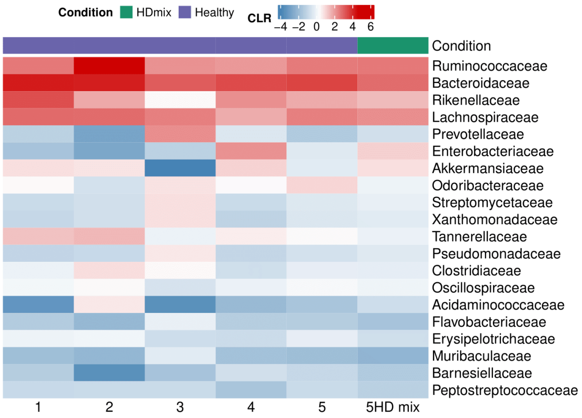


**c**

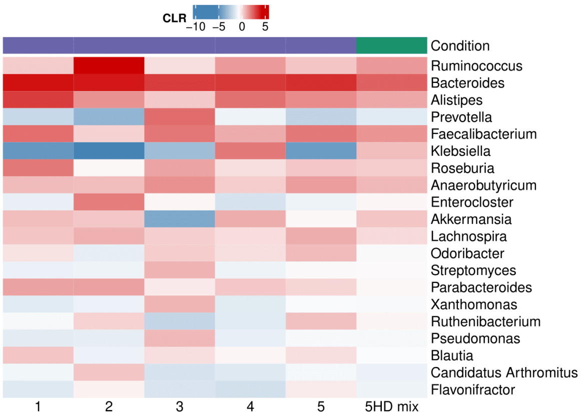


**d**

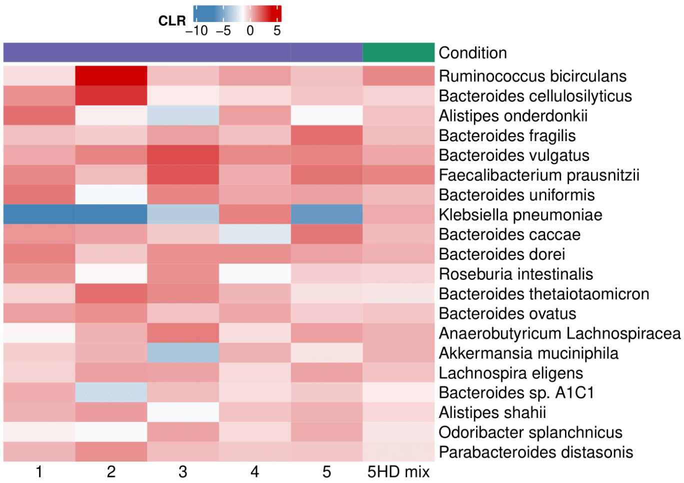


**e**

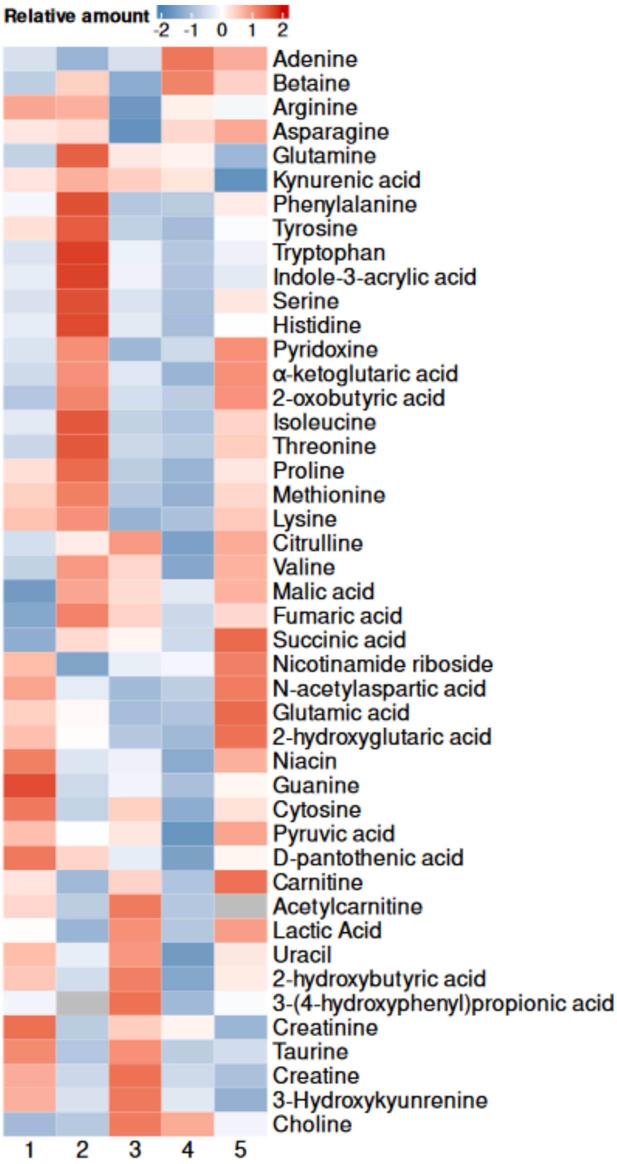

### Extended data figure 2

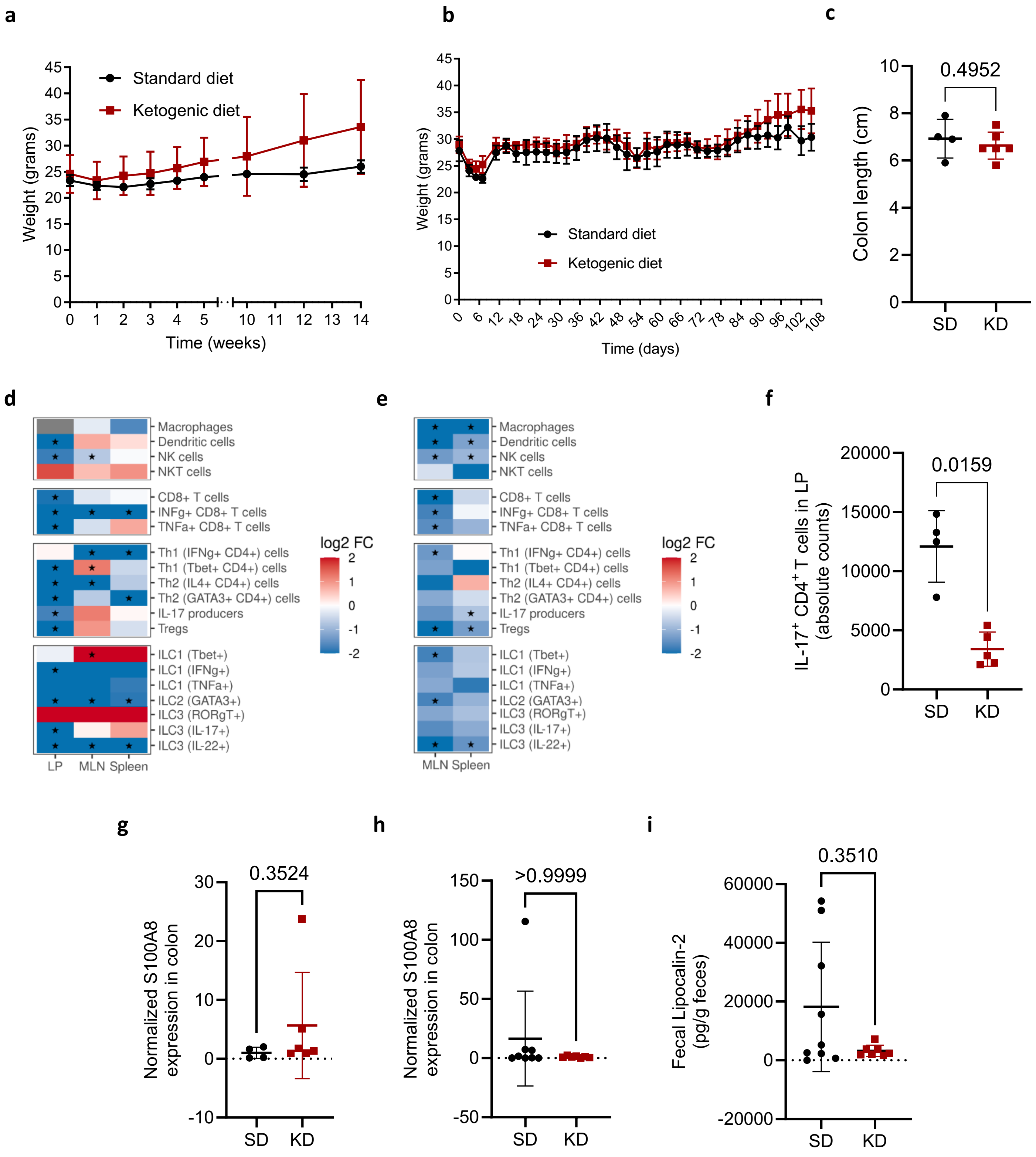

### Extended data figure 3

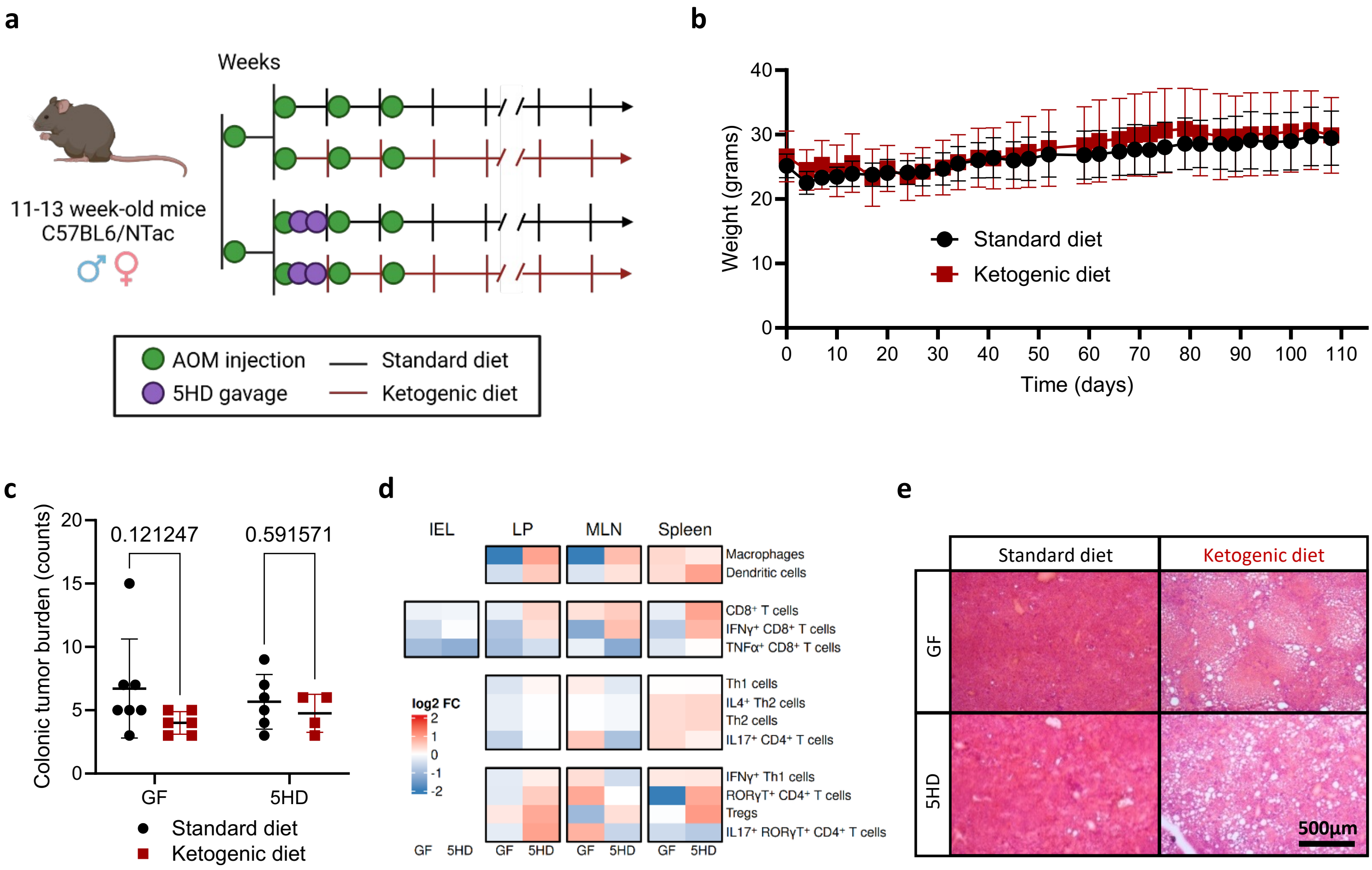

### Extended data figure 4

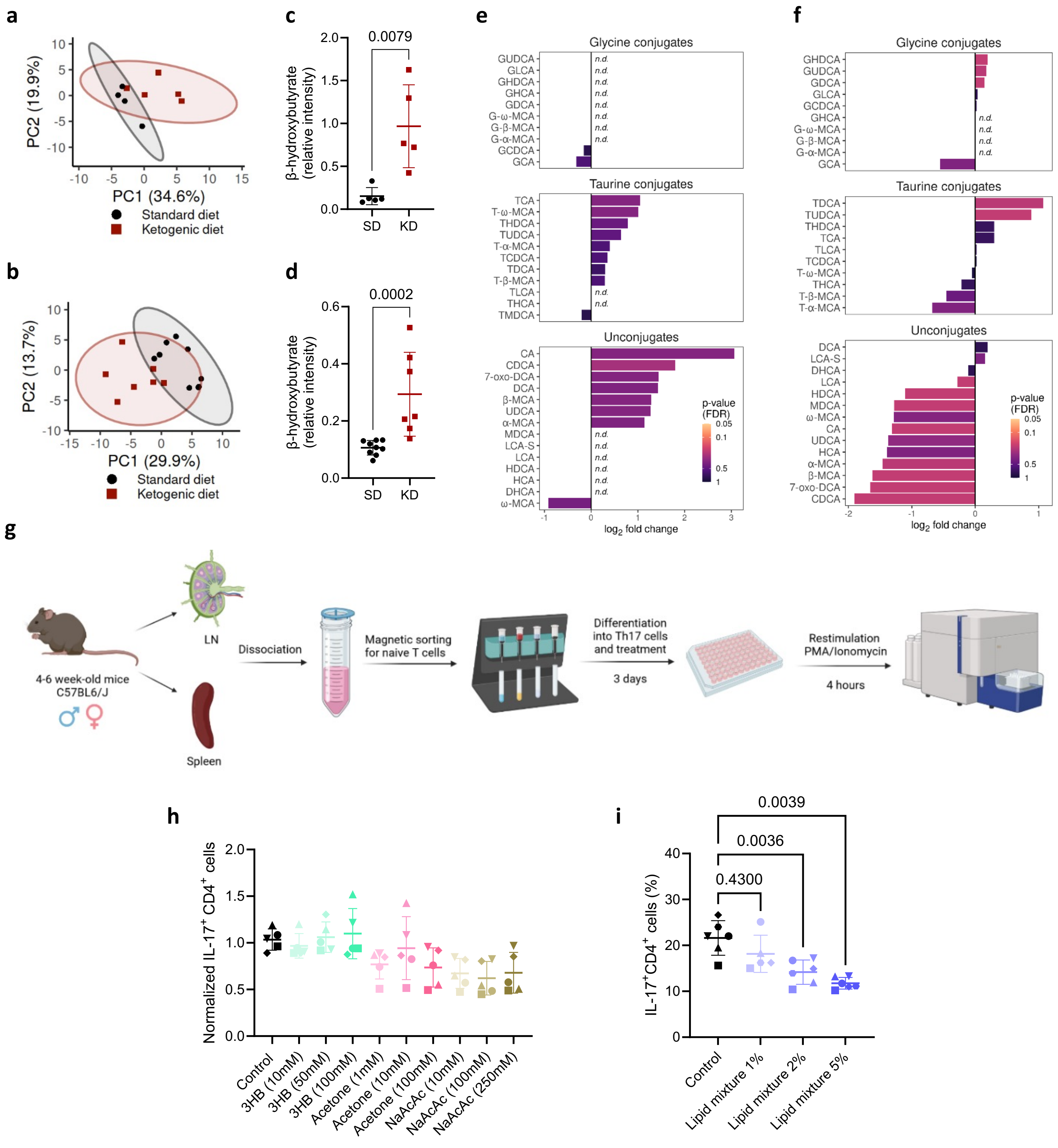

### Extended data figure 5

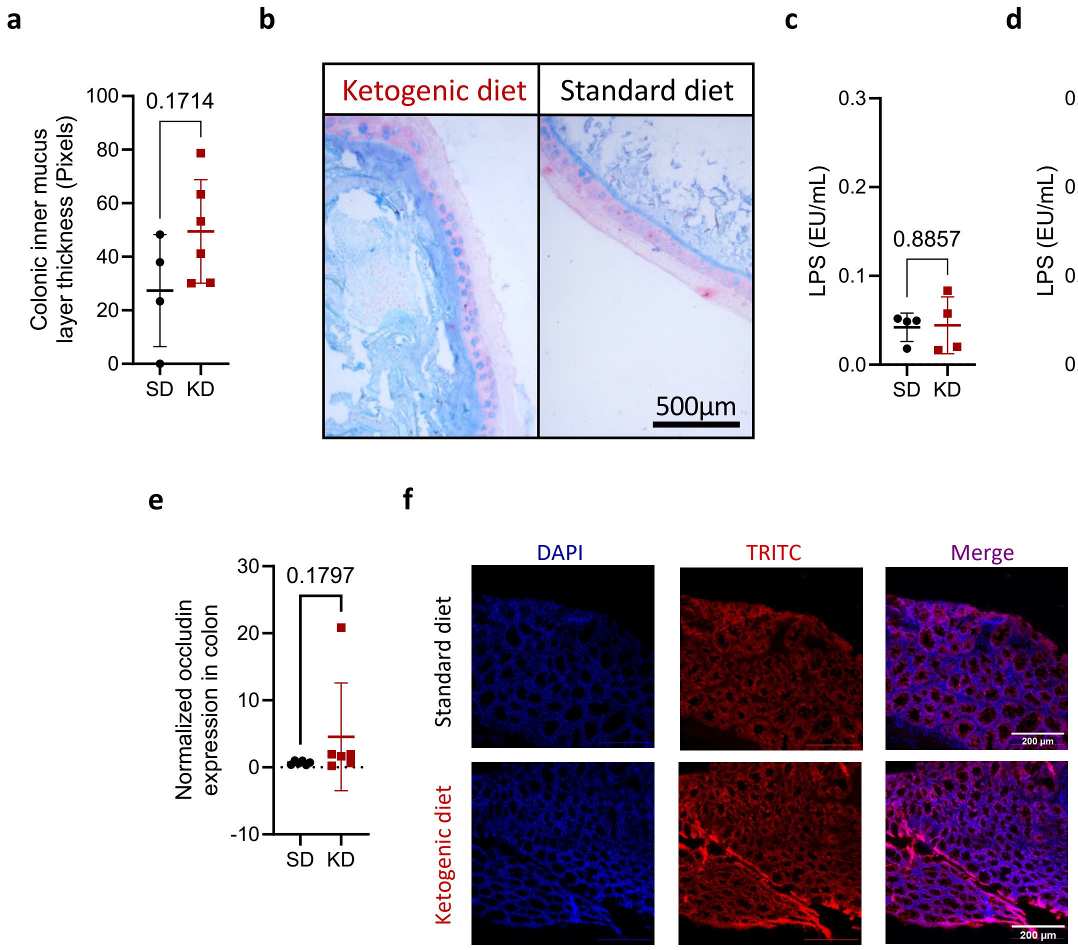

### Extended data figure 6

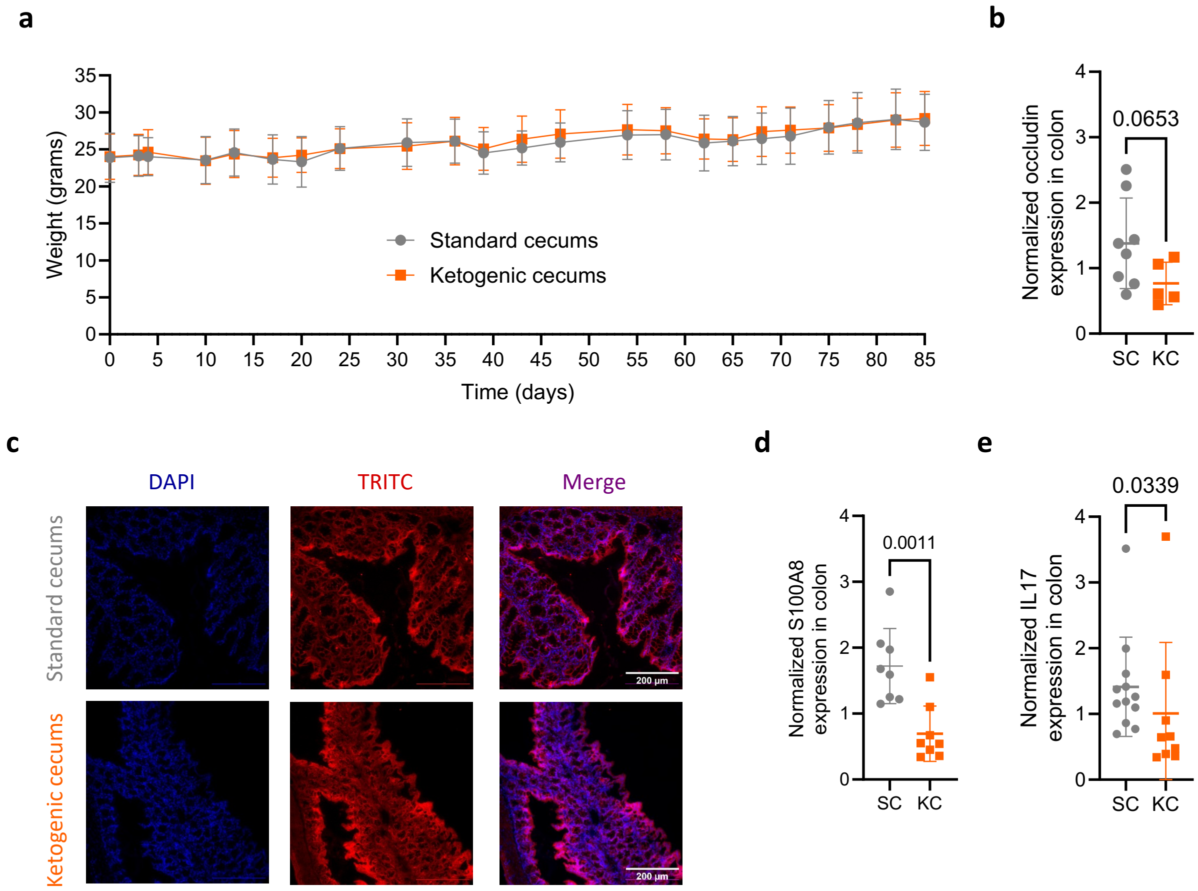

### Extended data figure 7

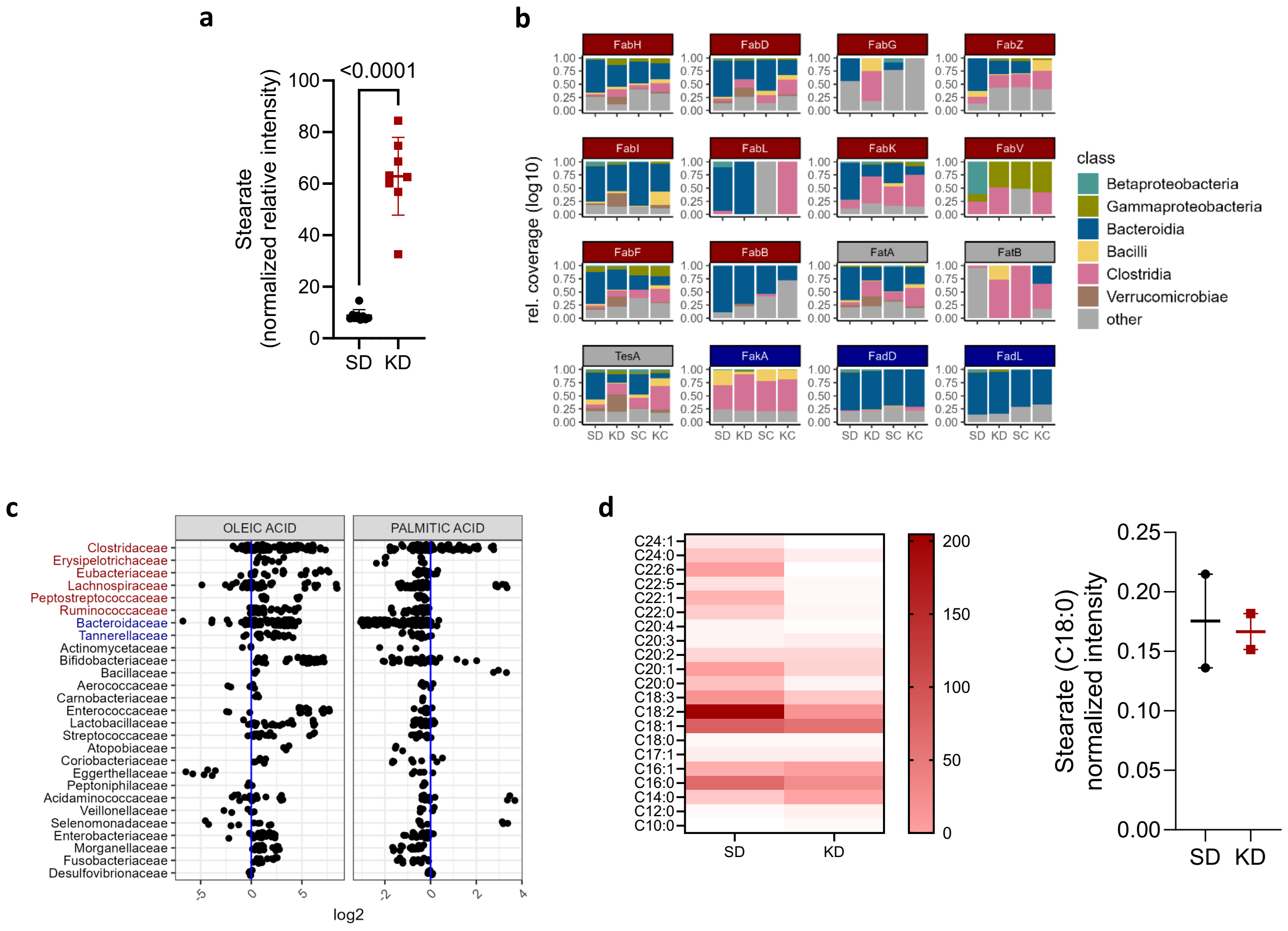

### Extended data figure 8

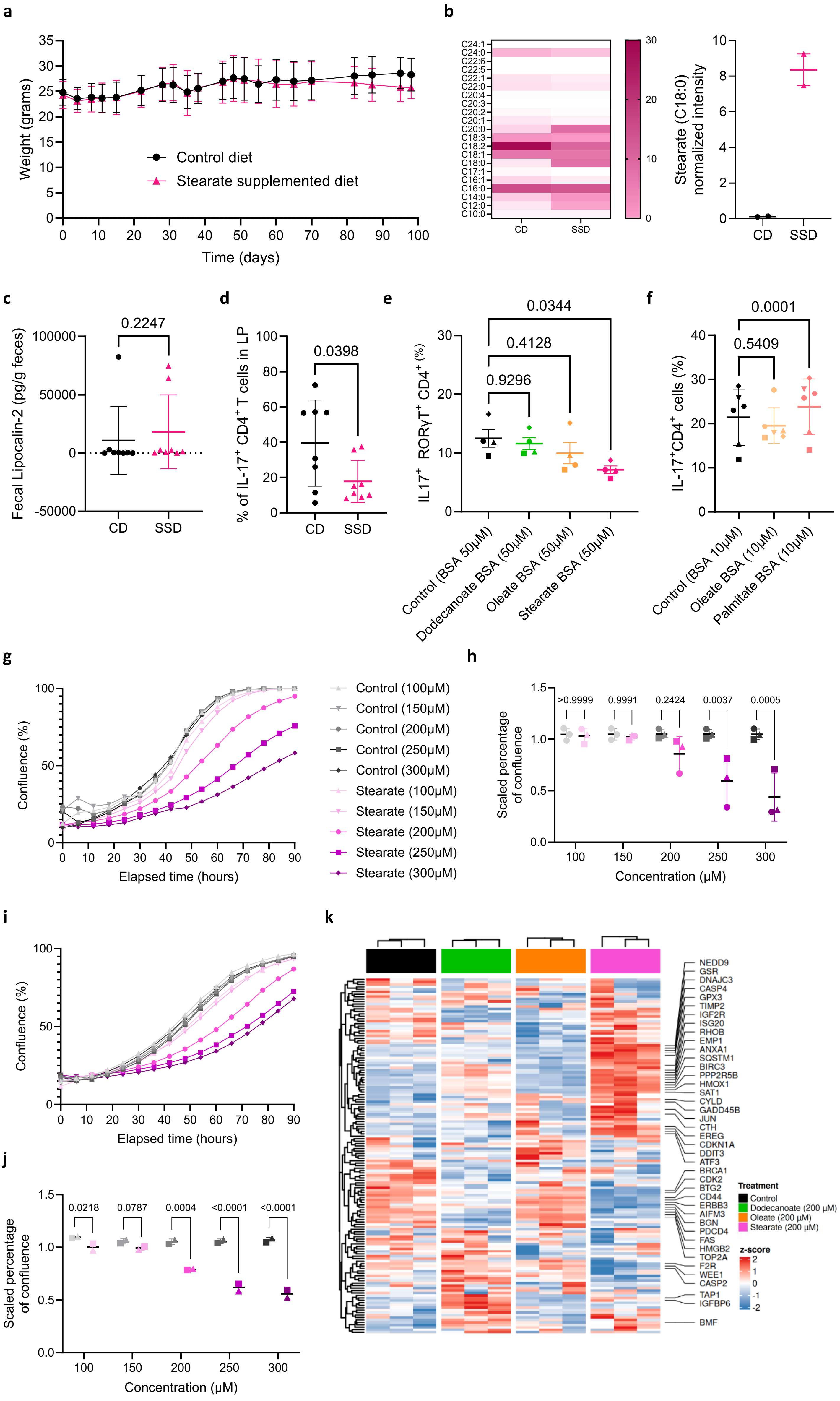

### Extended data figure 9

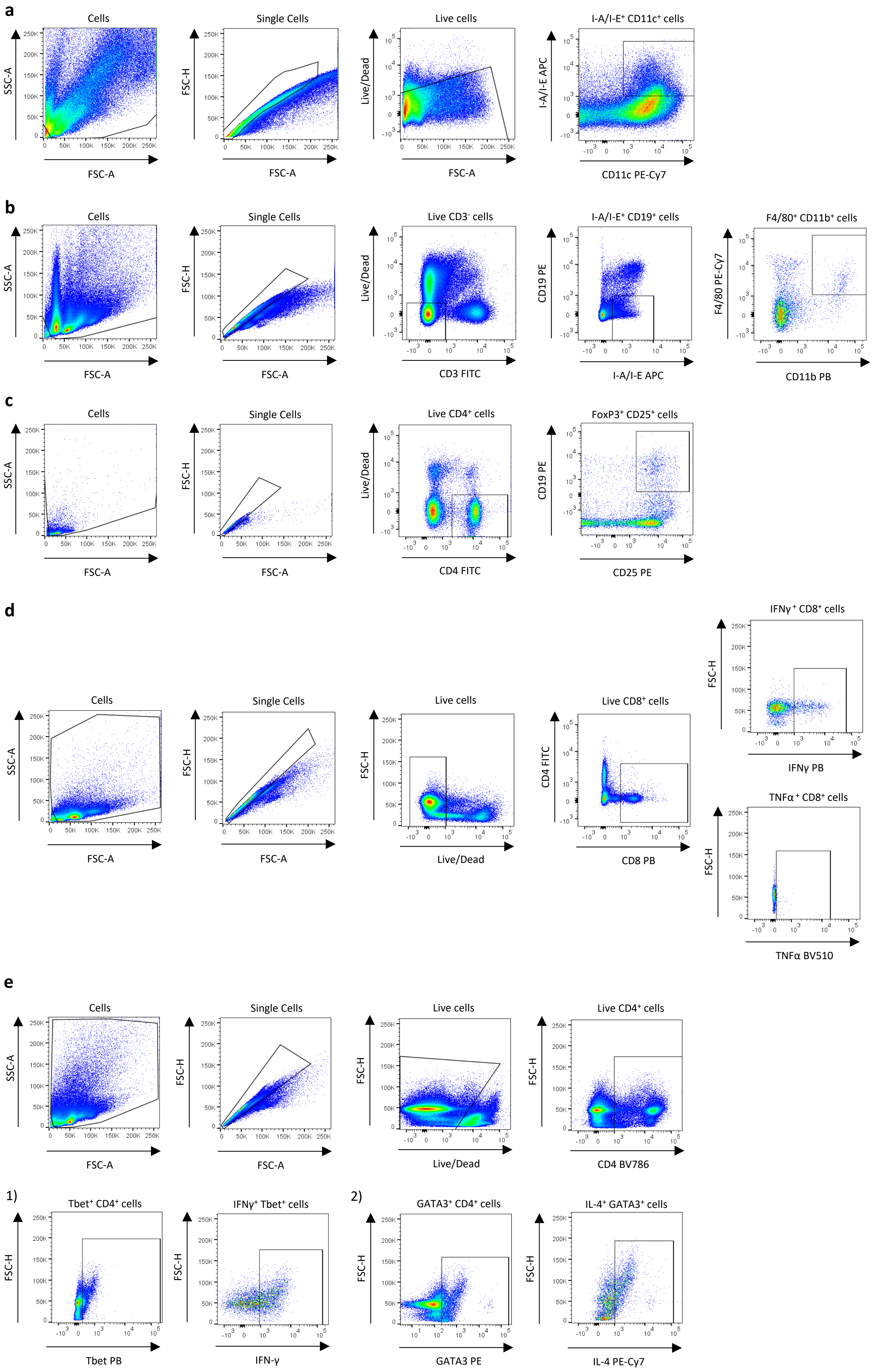

### Extended data figure 10

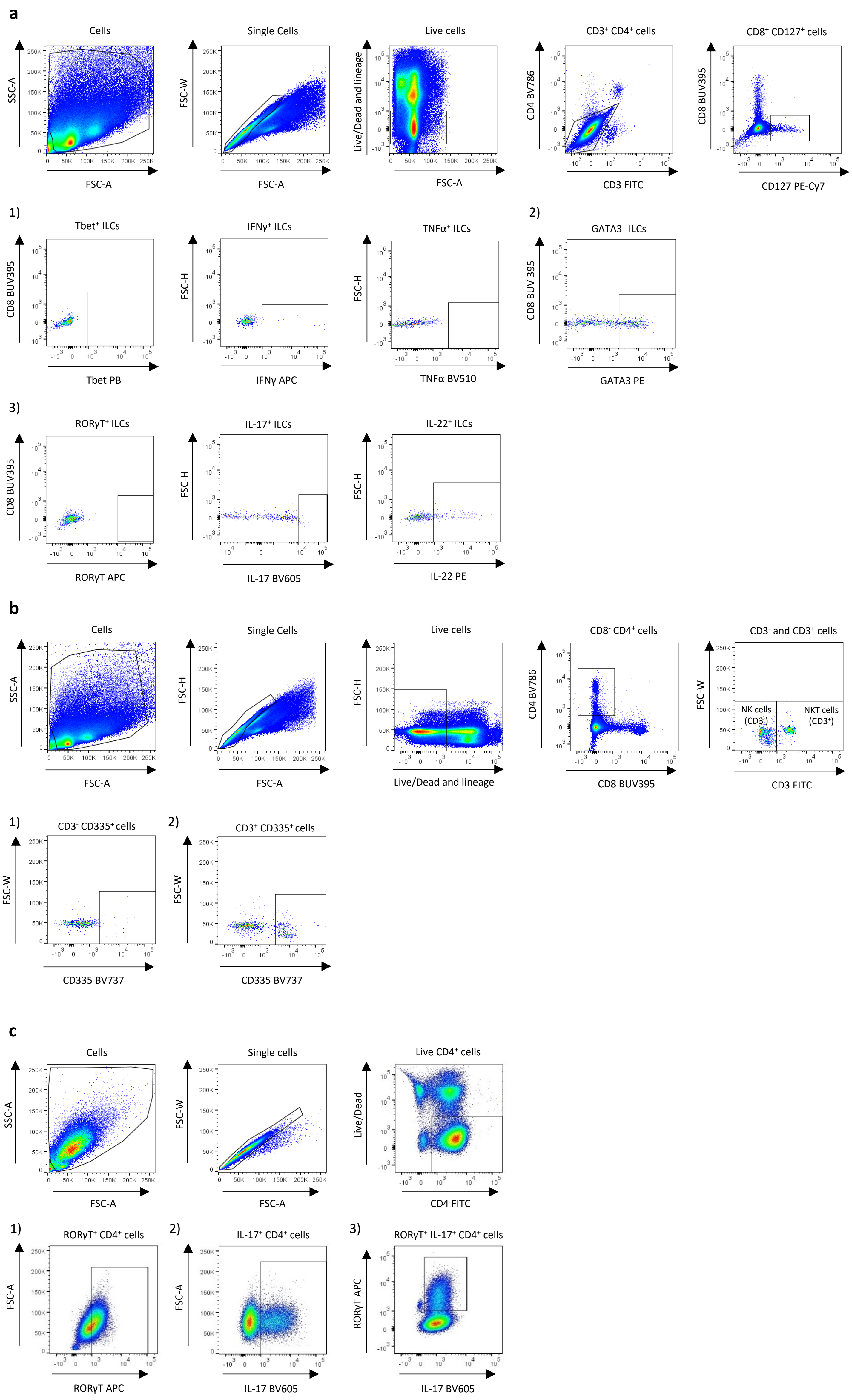
